# Conserved and unique transcriptional features of pharyngeal arches in the skate (*Leucoraja erinacea*) and evolution of the jaw

**DOI:** 10.1101/2021.01.12.426343

**Authors:** Christine Hirschberger, Victoria A. Sleight, Katharine E. Criswell, Stephen J. Clark, J. Andrew Gillis

## Abstract

The origin of the jaw is a long-standing problem in vertebrate evolutionary biology. Classical hypotheses of serial homology propose that the upper and lower jaw evolved through modifications of dorsal and ventral gill arch skeletal elements, respectively. If the jaw and gill arches are derived members of a primitive branchial series, we predict that they would share common developmental patterning mechanisms. Using candidate and RNAseq/differential gene expression analyses, we find broad conservation of dorsoventral patterning mechanisms within the developing mandibular, hyoid and gill arches of a cartilaginous fish, the skate (*Leucoraja erinacea*). Shared features include expression of genes encoding members of the ventralising BMP and endothelin signalling pathways and their effectors, the joint markers bapx1 and gdf5 and pro-chondrogenic transcription factors barx1 and gsc, and the dorsalising transcription factor pou3f3. Additionally, we find that mesenchymal expression of *eya1/six1* is an ancestral feature of the mandibular arch of jawed vertebrates, while differences in notch signalling distinguish the mandibular and gill arches in skate. Comparative transcriptomic analyses of mandibular and gill arch tissues reveal additional genes differentially expressed along the dorsoventral axis of the pharyngeal arches, including *scamp5* as a novel marker of the dorsal mandibular arch, as well as distinct transcriptional features of mandibular and gill arch muscle progenitors and developing gill buds. Taken together, our findings reveal conserved patterning mechanisms in the pharyngeal arches of jawed vertebrates, consistent with serial homology of their skeletal derivatives, as well as unique transcriptional features that may underpin distinct jaw and gill arch morphologies.

## Introduction

The jaw is an iconic example of anatomical innovation, and a uniting feature of the jawed vertebrate (gnathostome) crown group (Gans & Northcutt 1983; Mallatt, 1996; Northcutt, 2005). Over a century ago, the anatomist Karl Gegenbaur proposed a scenario of serial homology, whereby the upper and lower jaw arose through modifications of the dorsal and ventral elements of an anterior gill arch (Gegenbaur, 1878 - Fig. 1A). This hypothesis was based largely on the strikingly similar anatomical organization of the jaw and gill arches of cartilaginous fishes (sharks, skates and rays), and has since gained wide acceptance as a textbook scenario of jaw origin (Goodrich, 1930; de Beer, 1971; Romer, 1966; Carroll, 1988; though see Janvier, 1996 and Miyashita, 2015 for review and critical discussion of this hypothesis) (Fig. 1B). However, a series of transitional fossils showing the stepwise acquisition of the jaw along the gnathostome stem is lacking, and this gap in the fossil record has made it difficult to support or refute hypotheses of jaw-gill arch serial homology with palaeontological data. Elements of Gegenbaur’s hypothesis are, nevertheless, testable from a developmental perspective. Over the past several decades, concepts of serial homology have evolved to centre largely around the iterative deployment or sharing of conserved developmental mechanisms (e.g. Van Valen 1982; Roth, 1984; Wagner, 1989, 2007, 2014). If the parallel anatomical organisation of the gnathostome jaw and gill arch skeleton is a product of serial homology, we predict that these elements would be delineated by shared patterning mechanisms – and, conversely, that their anatomical divergence may be attributable to arch-specific variations on a core, conserved developmental programme.

**Figure 1:**
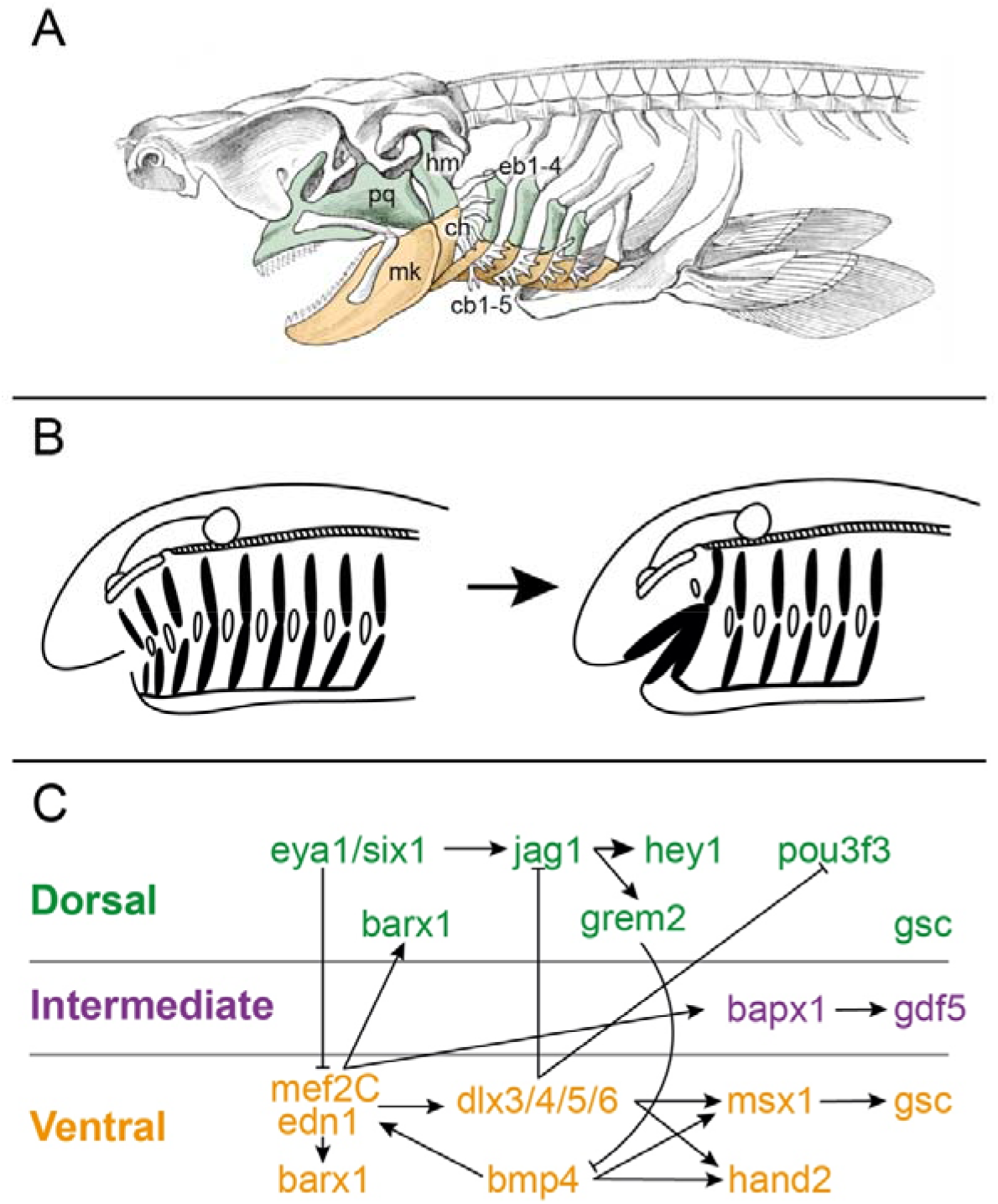
Anatomy, evolution and patterning of the pharyngeal endoskeleton. (A) Shark head skeleton illustrating an hypothesis of serial homology of the jaw and gill arch skeleton. Upper (dorsal) jaw, hyoid and gill arches are green, lower (ventral) jaw, hyoid and gill arches are orange (schematic modified from Owen, 1866). (B) Representative textbook scenario of jaw origin by transformation of an anterior gill arch (redrawn from Janvier, 1996 and references therein). (C) Signalling pathways and downstream effectors patterning the dorsoventral axis of the jaw, as established largely from studies in mouse and zebrafish (Redrawn after Cerny et al., 2010 and Medeiros & Crump, 2012). cb1-5, ceratobranchials 1-5; ch, ceratohyal; eb1-5, epibranchials 1-5; hm, hyomandibula; mk, Meckel’s cartilage; pq, palatoquadrate.

The endoskeletal elements of the jaw and gills develop from pharyngeal arches – transient, segmentally repeated columns of mesoderm and neural-crest-derived mesenchyme encased by epithelium in the embryonic vertebrate head (Graham, 2003). These embryonic tissues give rise to different elements of the craniofacial anatomy: head musculature forms from the pharyngeal arch core mesoderm, skeletal and connective tissue elements derive from neural crest and mesodermal mesenchyme, epidermal covering and sensory neurons derive from the ectodermal epithelium, and the inner lining of the pharynx and associated endocrine organs derive from the endoderm. In fishes, the first (mandibular) pharyngeal arch gives rise to the jaw skeleton, the second (hyoid) arch gives rise to a gill bearing arch that also functions, in some lineages, to suspend the jaw from the braincase, and a variable number of gill arches give rise to the skeletal support of the gills. Primitively, the skeletal derivatives of the pharyngeal arches were segmented, principally, dorsoventrally into the palatoquadrate and Meckel’s cartilage in the jaw, the hyomandibula and ceratohyal in the hyoid arch, and the epi- and ceratobranchial elements in the gill arches (de Beer, 1971; Janvier, 1996) (Fig 1A).

Studies in zebrafish and mouse have revealed a network of signalling interactions and transcription factors that are key to the development and patterning of the dorsal and ventral segments of the jaw in bony vertebrates (Fig 1C). Endothelin-1 (edn1) and bone morphogenetic protein 4 (bmp4) signalling from ventral mandibular arch epithelium and mesoderm promotes ventral expression of *dlx5/6, hand2* and *msx1/2* and imparts lower jaw identity (Ozeki et al., 2004, Miller et al., 2003; Clouthier et al., 1998; Beverdam et al., 2002; Depew et al., 2002; Yanagisawa et al., 2003; Alexander et al., 2011; Zuniga et al., 2011). Conversely, notch signalling (Barske et al., 2016; Zuniga et al., 2010) and *six1* expression (Tavares et al., 2017) repress transcription of *edn1*, and promote transcription of the upper jaw marker *pou3f3* (Jeong et al., 2008; Barske et al., 2020), thereby imparting upper jaw identity. Finally, the jaw joint is specified at the interface of these upper and lower jaw gene expression domains, with the presumptive joint marked by the expression of *bapx1/nkx3.2* (Miller et al., 2003; Lukas and Olsson, 2018) and *gdf5* (Miller et al., 2003), and flanked by expression of the pro-chondrogenic (and joint-repressing) transcription factor *barx1* (Nichols et al., 2013).

Taken together, these signalling interactions and transcription factors establish a dorsoventral (DV) code of combinatorial gene expression that confers axial identity on the mandibular and hyoid arch skeleton of bony fishes (Fig. 1C), though whether/which of these mechanisms were primitively shared between the mandibular, hyoid and gill arches of gnathostomes remains unclear. We, and others, have previously shown that nested expression of the *dlx* family of transcription factors, a key regulator of DV axial identity in the mandibular arch (Depew et al., 2002, 2005; Beverdam et al., 2002; Talbot et al., 2010), was primitively shared across all pharyngeal arches in gnathostomes (Gillis et al., 2013; Compagnucci et al., 2013), and that dorsal and ventral domains of *dlx* gene expression delineate the principal segments of the jaw and gill arch skeleton in a conserved manner in a chondrichthyan, the skate (*Leucoraja erinacea*) (Gillis et al., 2013). These findings are consistent with hypotheses of serial homology of the palatoquadrate/Meckel’s cartilage and epi-/ceratobranchial gill arch elements, respectively, although the degree of conservation or divergence of upstream signals and downstream effectors of this *“dlx* code” in the mandibular and gill arches has not been fully investigated.

To test the hypothesis that the jaw and gill arches are patterned by a shared transcriptional network, we have investigated the molecular development of the pharyngeal arches in the skate. This group has retained the primitive dorsoventrally segmented organization of the gnathostome pharyngeal endoskeleton (i.e. a jaw and gill arch skeleton that is segmented into prominent palatoquadrate/Meckel’s cartilage epi-/ceratobranchial elements, respectively – Mallatt, 1996; Gillis et al., 2012), and, through comparison with its sister group, the bony fishes, allows us to infer anatomical and developmental conditions in the last common ancestor of gnathostomes. Using a combination of candidate gene and comparative transcriptomic approaches, we find that the transcriptional network patterning the DV axis of the developing jaw in bony fishes is largely conserved and shared by the mandibular, hyoid and gill arches of skate, consistent with the hypothesis of jaw-gill arch serial homology. We further resolve dorsal mesenchymal expression of *six1* and *eya1* as a primitive and unique feature of the mandibular arch, we report *scamp5* as a novel marker of the dorsal territory of the mandibular arch, and we report transcriptional differences associated with progenitors of jaw and gill arch-specific musculature and gill primordia. Taken together, our findings point to a conserved gene regulatory network underlying the primitively shared organisation of the gnathostome mandibular, hyoid and gill arch skeleton, and highlight additional transcriptional features that correlate with the developmental and anatomical diversification of jaws and gill arches within gnathostomes.

## Results and Discussion

### Conservation of ventral gene expression patterns in the skate mandibular, hyoid and gill arches

In mouse (Kurihara et al., 1994; Clouthier et al., 1998; Ozeki et al., 2004) and in zebrafish (Miller et al., 2000; Kimmel et al., 2007), *edn1* is expressed in ventral and intermediate mandibular and hyoid arch epithelium, and this edn1 signal is transduced within the adjacent arch mesenchyme through its receptor, ednra, and its downstream effector mef2C (Nair et al., 2007; Miller et al., 2007; Sato et al., 2008). *bmp4* is similarly expressed in ventral arch epithelium in mouse (Liu et al., 2005) and in zebrafish (Alexander et al., 2011), where its ventral patterning function is restricted by intermediate expression of *grem2*, which encodes a secreted Bmp antagonist (Zuniga et al., 2011). Together, edn1 and bmp4 signalling promote ventral mesenchymal expression of *hand2* and *msx1*, and confer lower jaw identity (Thomas et al., 1998; Yanagisawa et al., 2003; Funato et al., 2016).

We carried out a series of mRNA *in situ* hybridisation (ISH) experiments to test for shared expression of ventral patterning factors in the pharyngeal arches of skate embryos. We found that *edn1* is expressed in the ventral/intermediate epithelium of the mandibular, hyoid and gill arches (Fig. 2A, B), while *ednra* is expressed throughout the mesenchyme of all pharyngeal arches (Fig. 2C, D). We additionally tested for expression of the gene encoding another endothelin receptor, *ednrb*. Although *ednrb* mutant mice have not been reported to exhibit craniofacial defects (reviewed by Pla and Larue, 2003), skate embryos exhibit shared expression of *ednrb* in ventral and intermediate mesenchyme across all pharyngeal arches (Fig. 2E, F), hinting at conservation within gnathostomes of a skeletal patterning function of *ednrb* that has so far only been described in a jawless fish, the lamprey (*Petromyzon marinus*) (Square et al., 2020). We also found shared expression of *mef2C* in the ventral/intermediate domain of all pharyngeal arches (Fig. 2G). It has been demonstrated that *mef2C* is a transcriptional target edn1 signalling in cranial neural crest-derived mesenchyme (Miller et al., 2007), and so our findings point to shared *edn1* signalling between epithelium and mesenchyme of all pharyngeal arches in skate.

**Figure 2:**
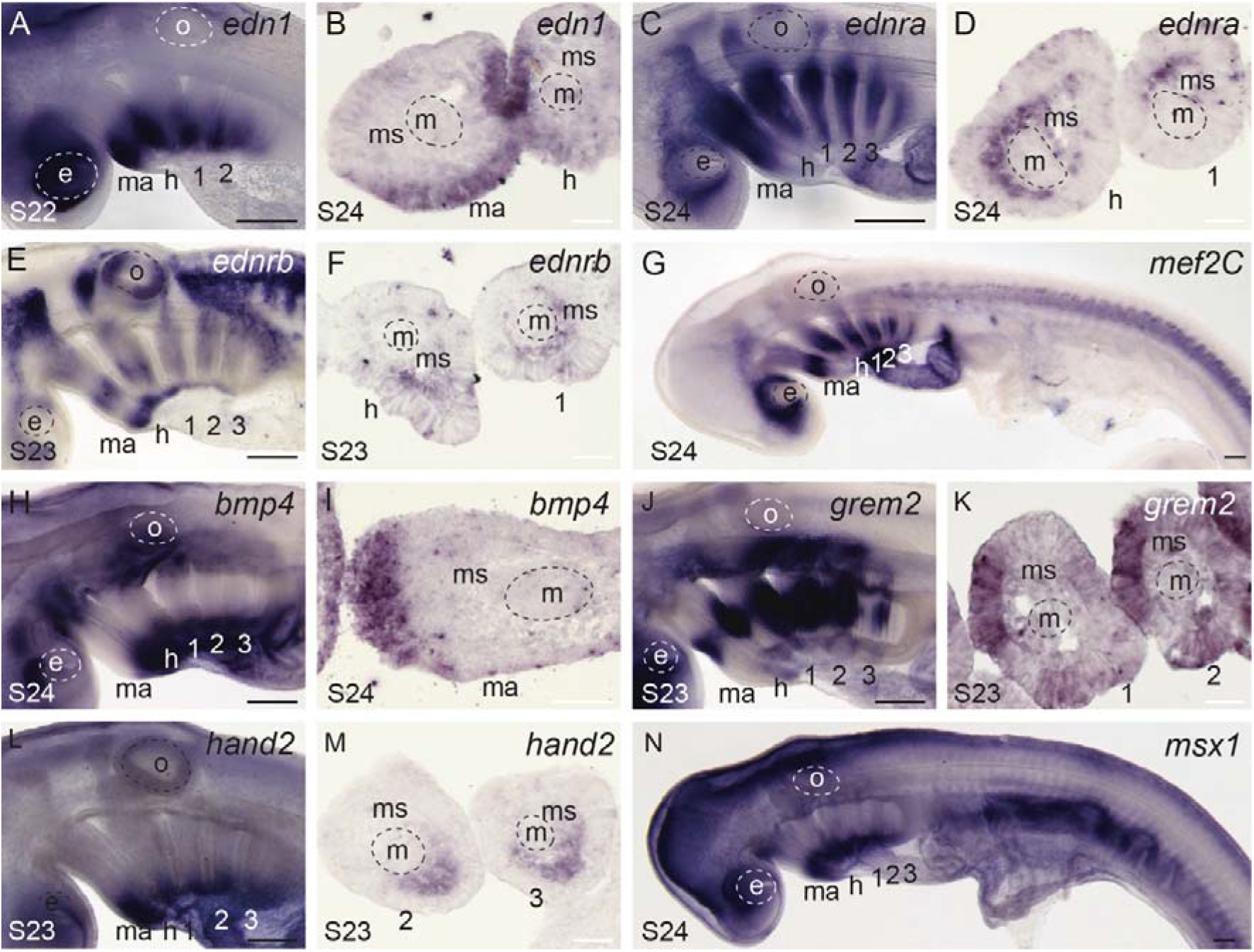
Conservation of ventral gene expression patterns in the skate mandibular, hyoid and gill arches. (A) At stage (S)22, *edn1* is expressed in the ventral domain of all pharyngeal arches, with transcripts localising to the (B) pharyngeal epithelium. (C) *ednra* is expressed along the entire DV axis of the pharyngeal arches, within (D) the mesenchyme. (E) *ednrb* is expressed in migrating neural crest streams, and also in distinct intermediate and ventral domains within (F) pharyngeal arch mesenchyme. (G) *mef2C* is expressed in the ventral and intermediate domains of all pharyngeal arches. (H) *bmp4* is expressed in ventral pharyngeal arch (I) epithelium, and (*J*) *grem2* is expressed in intermediate pharyngeal arch (K) epithelium. (L) *hand2* is expressed in the ventral (M) mesenchyme of each pharyngeal arch. (N) msx1 is expressed ventrally in all pharyngeal arches. 1, 2, 3: gill arches; e, eye; h, hyoid; m, mesoderm; ma, mandibular; ms, mesenchyme; o, otic vesicle. Scale bars: black 400um, white 25um.

We also tested for expression of bmp signalling components in skate pharyngeal arches, and found shared *bmp4* expression in the ventral epithelium of all arches (Fig. 2H, I). Dorsal to this *bmp4* domain, we observe shared intermediate/dorsal expression of *grem2* in the mandibular, hyoid and gill arch epithelium (Fig. 2J, K). Finally, we detect shared expression of *hand2* and *msx1* in the ventral mesenchyme of all pharyngeal arches (Fig. 2L-N). Taken together, our findings point to conservation of ventral pharyngeal arch patterning mechanisms between bony and cartilaginous fishes, and across the mandibular, hyoid and gill arches of the skate.

### Conserved and divergent dorsal expression of dorsal patterning genes in the skate mandibular, hyoid and gill arches

In mouse, *eya1* and *six1* function in craniofacial development (Xu et al., 1999; Laclef et al., 2003; Ozaki et al., 2004) and are co-expressed in the upper jaw primordium of the mandibular arch, where they inhibit expression of *edn1* and induce expression of the notch signalling component *Jag1* (Tavares et al., 2017). In zebrafish, *Jag1b* and *hey1* are expressed in the dorsal mesenchyme of the mandibular and hyoid arches and in pouch endoderm, while *notch2* is expressed more widely throughout the pharyngeal arches (Zuniga et al., 2010). Notch signalling through *Jag1b* and *hey1* confers upper jaw skeletal identity and restricts the expression of intermediate and ventral patterning genes, including *dlx3b/5a/6a, msxe, bapx1* and *barx1* (Zuniga et al., 2010; Barske et al., 2016). In zebrafish, *dlx2a* is also expressed throughout the DV mesenchyme axis of pharyngeal arches, and together with *dlx1a* functions to specify dorsal identity (Talbot et al., 2010) and, in mouse, to positively regulate the dorsal expression of another upper jaw marker within the arch mesenchyme, *pou3f3* (Jeong et al., 2008).

To test for conservation of dorsal patterning factors in the pharyngeal arches of the skate, we first characterised the expression of the transcription factors *eya1, six1* and *pou3f3* by ISH. We found that *six1* (Fig. 3A-C) and *eya1* (Fig. 3D-F) are both expressed broadly in the mandibular, hyoid, and gill arches in skate. However, while *six1* and *eya1* expression in the epithelium and mesodermal core is shared across the mandibular (Fig. 3B, E), hyoid and gill arches (Fig. 3C, F), mesenchymal expression of these factors is uniquely observed in the dorsal mandibular arch (Fig. 3B, E). Our findings are consistent with *six1* expression reported in mouse (Tavares et al., 2017) and chick (Fonseca et al., 2017), and point to an ancestral role for *eya1/six1* in patterning the upper jaw skeleton of gnathostomes. In contrast, *pou3f3* is expressed in the dorsal mesenchyme of the mandibular, hyoid, and gill arches (Fig. 3G-I), indicating a likely shared role in dorsal patterning across all pharyngeal arches.

**Figure 3:**
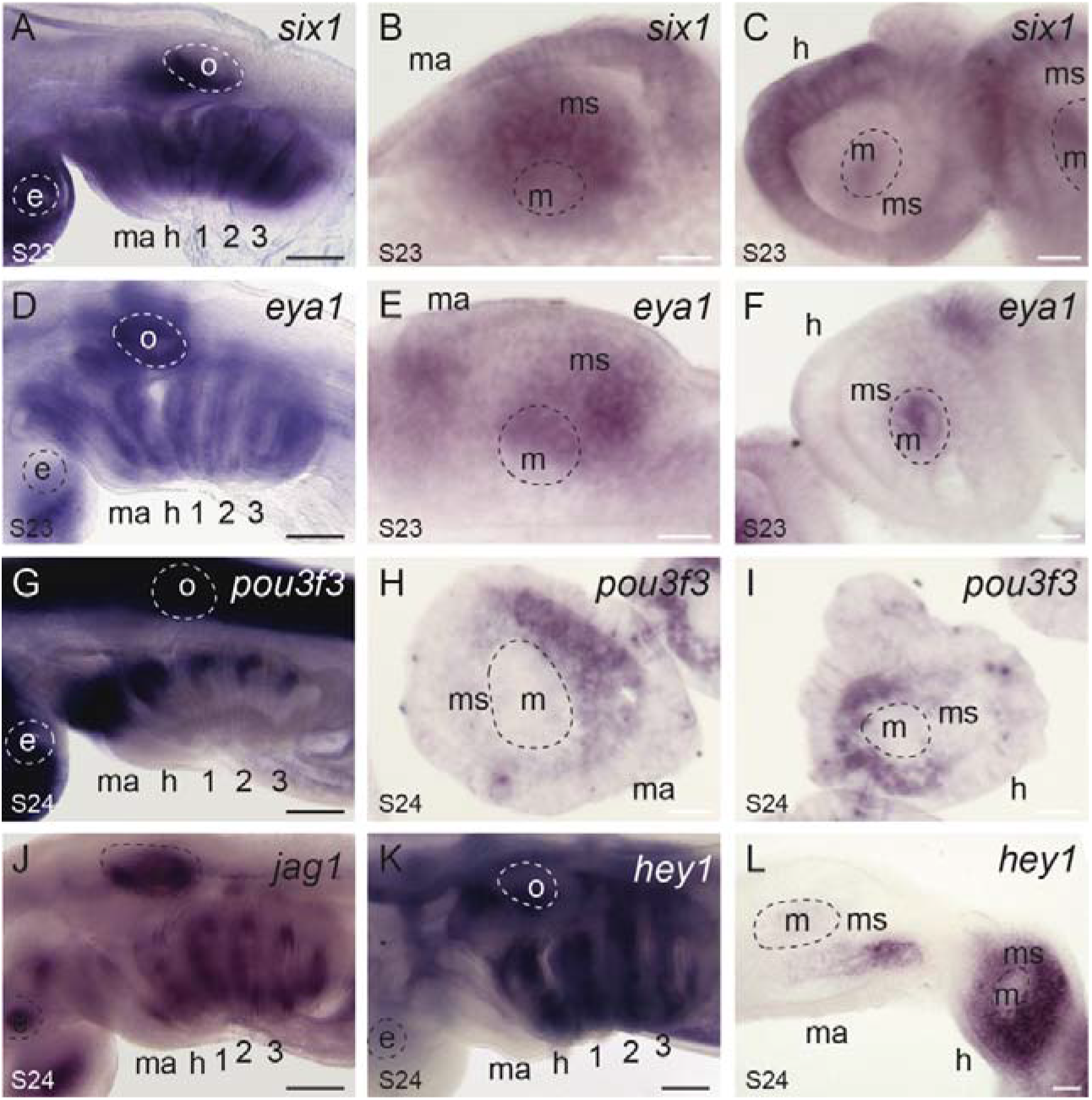
Conserved and divergent dorsal gene expression patterns in the skate mandibular, hyoid and gill arches. (A) *six1* is expressed in the (B) mesenchyme, core mesoderm and epithelium of the mandibular arch, and in the (C) the core mesoderm and epithelium of the hyoid and gill arches. Similarly, (D) *eya1* is expressed in the (E) mesenchyme, core mesoderm and epithelium of the mandibular arch, and in the (F) the core mesoderm and epithelium of the hyoid and gill arches. (G) *pou3f3* is expressed in the dorsal mesenchyme of the (H) mandibular, (I) hyoid and gill arches. (*J*)*jag1* is expressed in the mandibular, hyoid and gill arches, though (K) the notch signalling readout *hey1* is expressed (L) in a very restricted pattern within the mandibular arch mesenchyme, but broadly throughout the hyoid and gill arch mesenchyme. 1, 2, 3: gill arches; e, eye; h, hyoid; m, mesoderm; ma, mandibular; ms, mesenchyme; o, otic vesicle. Scale bars: black 400um, white 25um.

We next tested for expression of genes encoding the notch signalling components jag1 and hey1. We observe *Jag1* expression in the hyoid and gill arches of skate, but not in the mandibular arch (with the exception of very restricted expression in the posterior mandibular arch epithelium – Fig. 3J). In line with this, we also detect strong expression of *hey1* (a notch signalling readout) throughout the mesenchyme of the hyoid and gill arches, but only very restricted expression within a subdomain of the posterior mandibular arch mesenchyme (Fig. 3K, L; Fig. S1A-F). These observations differ from patterns previously reported in zebrafish, both in terms of DV extent of expression (i.e. expression along the entire DV extent of the arch in skate, as opposed to the dorsal localisation seen in zebrafish), and the near exclusion of mesenchymal *hey1* expression from the mandibular arch in skate. It is possible that the upper jaw patterning function of jag1 signalling is an ancestral feature of the gnathostome mandibular arch that has been lost or reduced in skate, or that this mechanism is a derived feature of bony fishes. Gene expression data for notch signalling components in the pharyngeal arches of cyclostomes (lampreys and hagfishes) are needed to resolve this.

### Conservation of joint gene expression patterns in the skate mandibular, hyoid and gill arches

In bony fishes, the jaw joint is specified by expression of genes encoding the transcription factor bapx1 and the secreted signalling molecule gdf5, and is flanked by expression of genes encoding the pro-chondrogenic transcription factors barx1 and gsc (Miller et al., 2003; Nichols et al., 2013; Wilson and Tucker, 2004; Newman et al., 1997, Lukas and Olsson, 2018; Tucker et al., 2004; Trumpp et al, 1999). In skate, we observe shared mesenchymal expression of *barx1* (Fig. 4A, B) and *gsc* (Fig. 4C) in the dorsal and ventral domains of the mandibular, hyoid and gill arches, and later, complementary mesenchymal expression of *gdf5* (Fig. 4D,E) and *bapx1* (Fig. 4F,G) in the intermediate region of all arches. These expression patterns are consistent with conservation of the pro-chondrogenic function of *barx1* and *gsc*, and the joint patterning function of *bapx1* and *gdf5*, in cartilaginous fishes.

**Figure 4:**
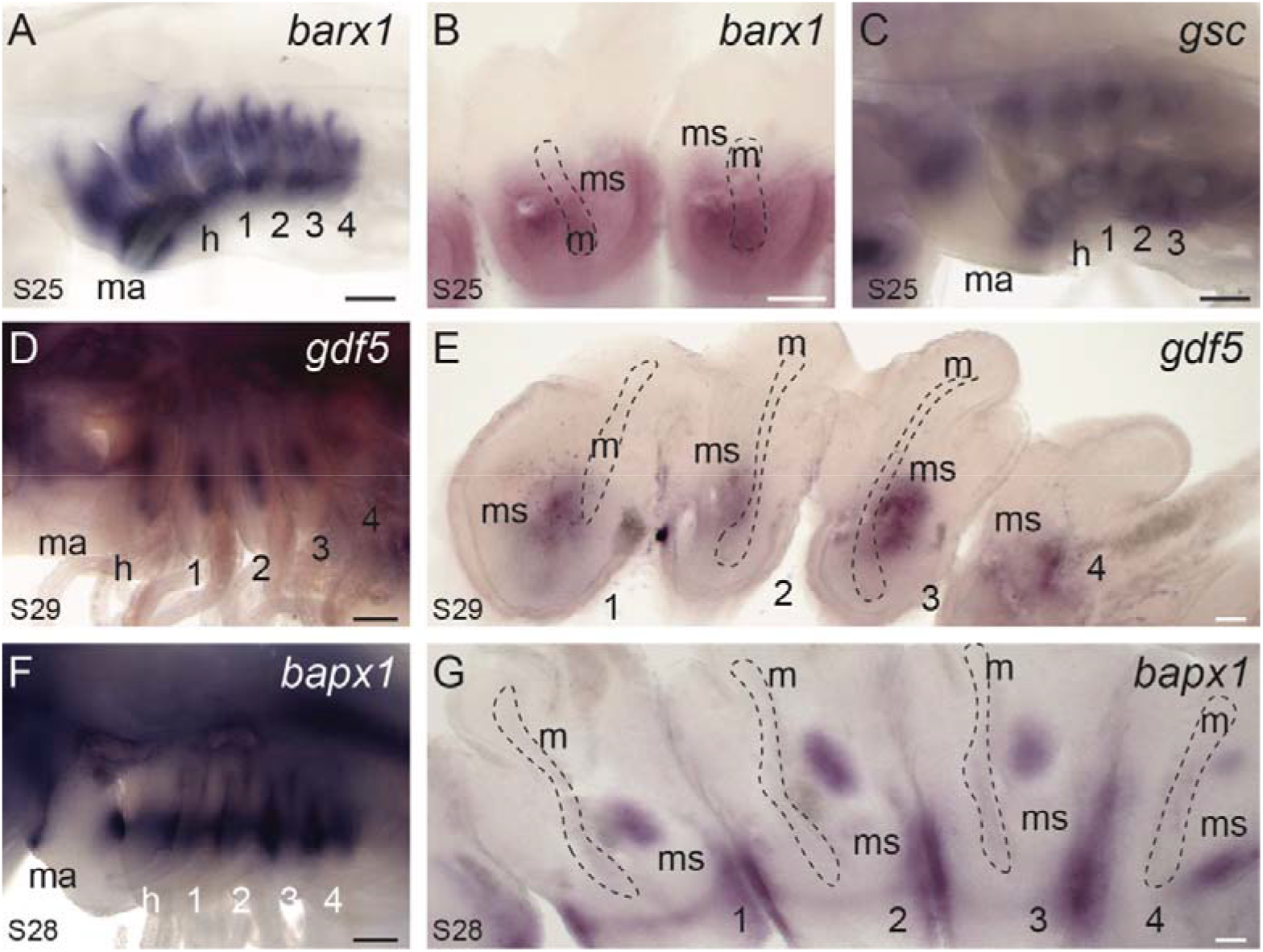
Conserved expression of joint markers and pro-chondrogenic transcription factors in the skate mandibular, hyoid and gill arches. (A) *barx1* is expressed in dorsal and ventral (B) mesenchyme across all pharyngeal arches, in a pattern that flanks the presumptive joint domain. (C) *gsc* is also expressed in dorsal and ventral domains of all pharyngeal arches, excluding the intermediate, presumptive joint domains. (D) *gdf5* is subsequently expressed in the intermediate (E) mesenchyme of all pharyngeal arches. (F) *bapx1* is expressed in the intermediate (G) mesenchyme and epithelium of all pharyngeal arches. 1, 2, 3: gill arches; e, eye; h, hyoid; m, mesoderm; ma, mandibular; ms, mesenchyme; o, otic vesicle. Scale bars: black 400um, white 25um.

A previous study of axial patterning gene expression in the pharyngeal arches of the jawless lamprey reported broad conservation of *dlx, hand* and *msx* expression across all pharyngeal arches, but a conspicuous absence of ł>αpxand *gdf* expression in the intermediate region of the first arch. These observations led to the suggestion that co-option of these joint patterning factors to the intermediate region of the mandibular arch, on top of a preexisting and deeply conserved DV patterning programme, was key to the evolutionary origin of the jaw (Cerny et al., 2010). Our findings are consistent with acquisition of intermediate *bapx1* and *gdf5* expression as a key step in the origin of the jaw joint, but suggest that this developmental mechanism was not primitively mandibular arch-specific, but rather a conserved mechanism specifying joint fate in the skeleton of the mandibular, hyoid and gill arches of gnathostomes.

### Comparative transcriptomics reveals additional mandibular and gill arch DV patterning genes

In an attempt to discover additional factors involved in DV patterning of the pharyngeal skeleton, we performed a comparative transcriptomic and differential gene expression analysis of upper and lower jaw and gill arch progenitors from skate embryos from S23-26. It is during these stages that DV axial identity is established within skate pharyngeal arches, as evidenced by nested expression within pharyngeal arches of the *dlx* family of transcription factors (Gillis et al., 2013), and by expression of the known axial patterning candidate genes characterised above. We manually dissected dorsal and ventral domains of the mandibular and first gill arch of S23/24 and S25/26 skate embryos (based on morphological landmarks correlating with dorsal and ventral *Dlx* code expression, after Gillis et al., 2013) (Fig. 5A), and performed RNA extraction, library preparation and RNAseq for each half-arch. After *de novo* transcriptome assembly, we conducted within-arch comparisons of gene expression levels between dorsal and ventral domains of the mandibular and first gill arch, and across-arch comparisons of gene expression levels between dorsal mandibular and dorsal gill arch domains, and between ventral mandibular and ventral gill arch gill arch domains (Fig. 5B-E, Fig. S1E-H).

**Figure 5:**
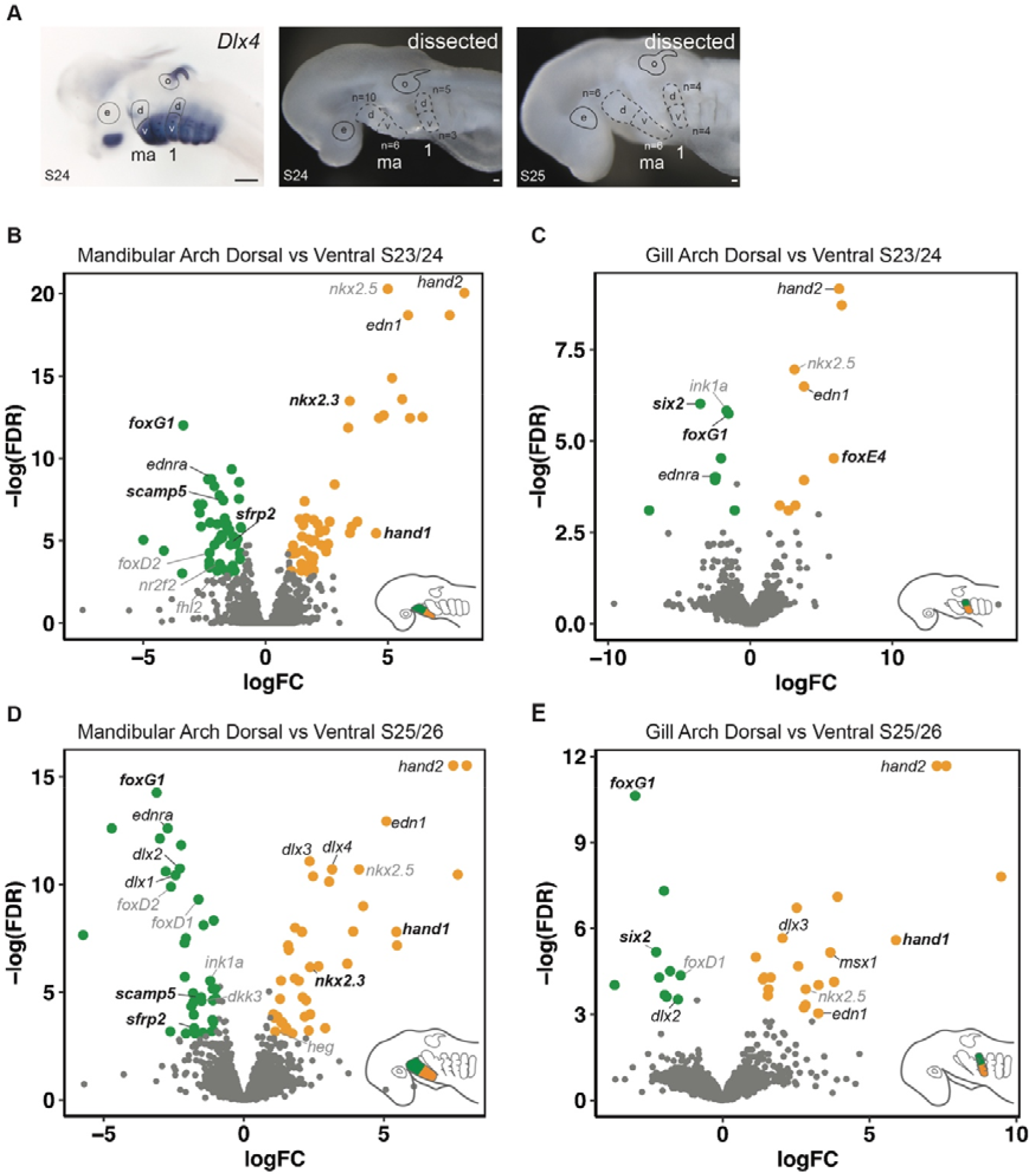
*De novo* transcriptome and differential gene expression analysis of dorsal and ventral domains of skate pharyngeal arches. (A) Demarcation of dorsal and ventral domains of the mandibular and gill arch based on previously published *Dlx* gene expression (Gillis et al., 2013). Dorsal and ventral domains of the mandibular and first gill arch were collected by manual dissection from skate embryos at S23/24 and S25/26. Volcano plots illustrate genes that are significantly differentially expressed within the dorsal and ventral domains of the (B) mandibular arch at S23/24, (C) the first gill arch at S23/24, (D) the mandibular arch at S25/26 and (E) the first gill arch at S25/26. Genes with established roles in pharyngeal arch axial patterning are in simple italics, additional genes for which we provide *in situ* validation are in bold italics, and additional factors highlighted by our analysis but not validated by mRNA *in situ* hybridisation are in grey italics. 1, first gill arch; *d*, dorsal; *e, eye; ma*, mandibular arch; *o*, otic vesicle, *v*, ventral. Scale bars: black 400um, white 25um.

We identified a number of transcripts as differentially expressed, defined as greater than a 2-fold change between tissue types with an adjusted P-value less than 0.05 (log2-fold changes [log2FC] > 1, P-value adjusted using Benjamin-Hochberd method < 0.05), within and between arch types at S23-24 and S25-26 (Table S2). Our ability to identify differentially expressed transcripts within and between arches using this approach was corroborated by the correct identification of known or expected genes within the appropriate spatial territory – e.g. *hand2, edn1* and *dlx3/4* were identified as differentially expressed within ventral territories (Fig. 5B, D), and *otx2* and *hox* genes were identified as differentially expressed within the mandibular and gill arch territories, respectively (Fig. S1E-H). To further biologically validate some of the findings of our analysis, we selected up to eight of the topmost differentially expressed transcription factors or signalling pathway components per comparison (excluding those already queried by our candidate gene approach or those with well-known functions in axial patterning of the pharyngeal skeleton), and attempted to clone fragments for *in situ* gene expressions analysis (Table S3 – complete lists of differentially expressed transcripts from each comparison are provided in Tables S4-S11). Out of 37 uniquely identified genes, we generated riboprobes for an additional 16 candidates, and we tested spatial expression of these candidates by mRNA *in situ* hybridisation.

We observed *foxG1* expression in the dorsal domains of the mandibular, hyoid and gill arches in skate (Fig. 6A). In mouse, *foxG1* functions in the morphogenesis of the forebrain (Tao and Lai, 1992; Dou et al., 1999; Hanashima et al., 2002), and more recently, has been shown to play a role in neurocranial and pharyngeal skeletal development (Compagnucci & Depew, 2020). In skate, we find that *foxG1* is expressed in the dorsal epithelium and dorsal core mesoderm of each pharyngeal arch (Fig. 6A, B), pointing to a likely ancestral role of this gene in regulating polarity of pharyngeal arch derivatives in gnathostomes, and a role in development of dorsal arch mesodermal derivatives in skate.

**Figure 6:**
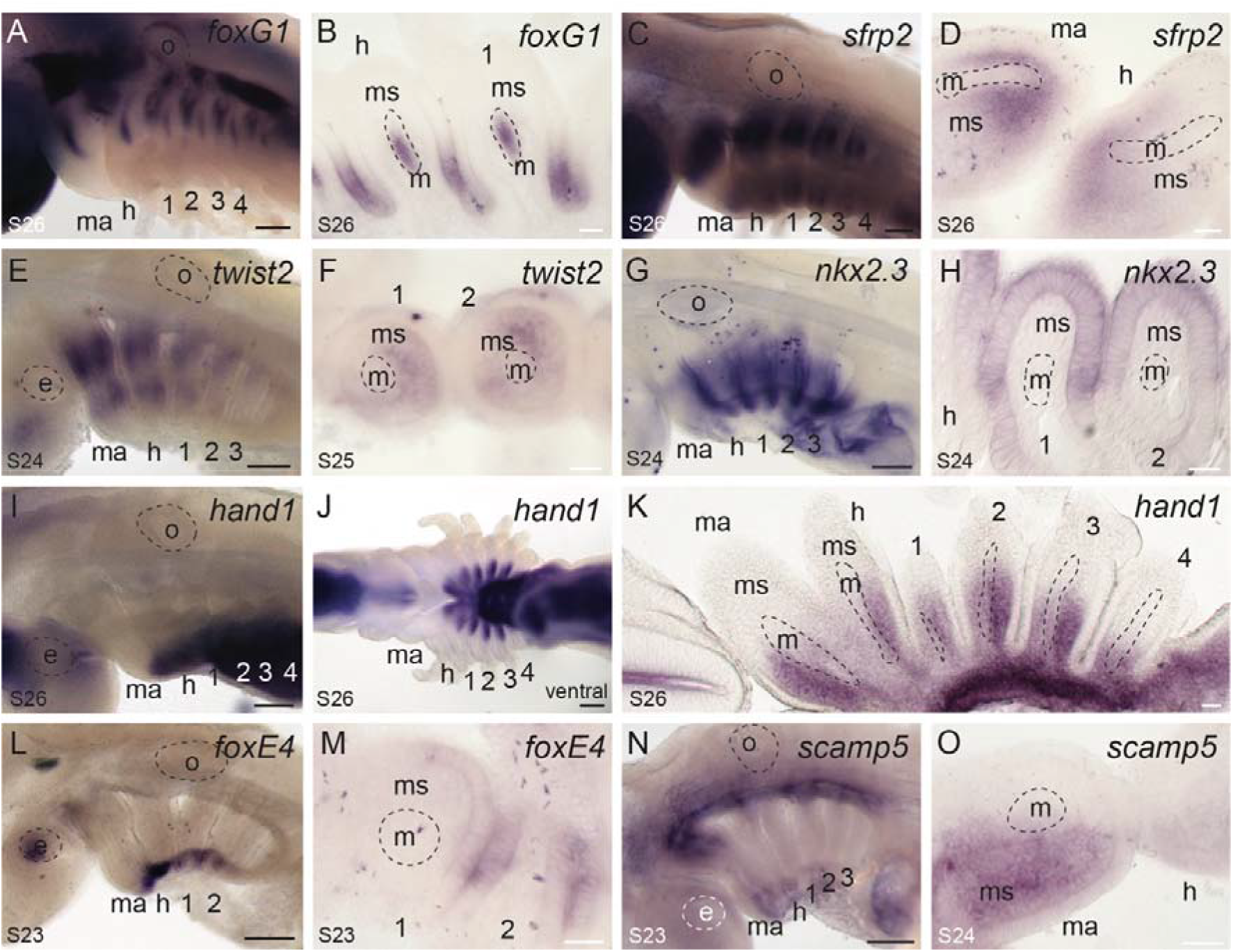
Additional genes exhibiting polarised expression along the DV axis of skate pharyngeal arches. (A) *foxG1* is expressed in dorsal (B) pharyngeal arch epithelium and core mesoderm of skate mandibular, hyoid and gill arches. (C) *sfrp2* is expressed in dorsal and ventral (D) mesenchyme of each pharyngeal arch. Similarly, (E) *twist2* is expressed in dorsal and ventral (F) mesenchyme of each pharyngeal arch. (G) *nkx2.3* is expressed in the ventral and intermediate (H) epithelium of each pharyngeal arch. (I) *hand1* transcripts localise to the (j) ventral (L) mesenchyme of each pharyngeal arch. (*L*)*foxE4* is expressed in the ventral extreme of the pharyngeal region, (M) with transcripts localising to the epithelium. (N) *scamp5* is expressed in the dorsal (0) mesenchyme of the mandibular arch. 1, 2, 3, 4: gill arches 1-4; e, eye; h, hyoid; m, mandibular arch; me, mesoderm; ms, mesenchyme; o, otic vesicle. Scale bars: black 400um, white 25um.

We additionally found that *sfrp2* (Fig. 6C) and *twist2* (Fig. 6E) are expressed in a discontiguous pattern, in the dorsal and ventral domains of skate pharyngeal arches. In chick, *sfrp2* is expressed in migrating cranial neural crest cells (Terry et al., 2000), while in mouse, it is expressed in the mesenchyme of the maxillary and mandibular domains of the mandibular arch (Leimeister et al., 1998). *sfrp2* is also expressed in the pharyngeal arches in zebrafish (Tendeng & Houart, 2006), where RNAseq experiments found it to be enriched in cranial neural crest cells of the dorsal mandibular and hyoid arches (Askary et al., 2017). However, wholemount fluorescent *in situ* hybridisation in zebrafish detected *sfrp2* expression only in the dorsal mesoderm, and TALEN and CRISPR induced early frameshift mutations in this gene did not lead to any observable skeletal craniofacial phenotypes (Askary et al., 2017). *twist2* is a basic helix-loop-helix transcription factor that is expressed in the dermis, cranial mesenchyme, pharyngeal arches and tongue of the mouse (Li et al., 1995), and in the mesenchyme of the mandibular and hyoid arches in chick (Scaal et al., 2001). Human nonsense mutations in *twist2* are linked to Setleis syndrome, a focal facial dermal dysplasia, and *twist2* knockout mice exhibit a similar facial phenotype (Tukel et al., 2010). In skate, we observed mesenchymal expression of both *sfrp2* (Fig. 6C,D) and *twist2* (Fig. 6E,F) in the dorsal and ventral mesenchyme of all pharyngeal arches, in patterns reminiscent of the pro-chondrogenic genes *barx1* and *gsc*, suggesting a possible role for these genes in the regulation of chondrogenesis.

Among genes with predicted expression in ventral pharyngeal arch territories, we found shared ventral expression of *nkx2.3, foxE4* and *hand1* across all pharyngeal arches in skate. *nkx2.3* is expressed in the endodermal lining of the pharynx in frog, mouse and zebrafish (Evans et al., 1995; Biben et al., 2004; Lee et al., 1996), and in skate, we find conservation of this pharyngeal endodermal expression (though with ventral endodermal localisation of *nkx2.3* transcripts at S24 - Fig. 6G, H). In mouse, *hand1* functions in cardiac morphogenesis (Srivastava et al., 1995; Riley et al., 1998), but is also expressed in the ventral mesenchyme of the pharyngeal arches (Clouthier et al., 2000). Targeted deletion of *hand1* alone does not result in craniofacial defects, though ablation of *hand1* on a *hand2* heterozygous background results in ventral midline defects within the jaw skeleton, suggesting a dosage dependent role for *hand* genes in mandibular skeletal patterning (Barbosa et al., 2007). Skate *hand1* is expressed in the ventral mesenchyme of each pharyngeal arch (Fig. 6I, J, K), in a pattern largely overlapping with the ventral mesenchymal expression of *hand2*, consistent with an ancestral combinatorial role for *Hand* genes patterning the ventral pharyngeal arch skeleton of gnathostomes. Finally, *foxE4* is expressed in the pharyngeal endoderm of non-teleost ray-finned fishes (Minarik et al., 2017), and in the endostyle (an endodermally-derived secretory organ and putative evolutionary antecendent of the thyroid gland) in non-vertebrate chordates (Yu et al., 2002; Hiruta et al., 2005). In skate, *foxE4* expression is conserved in ventral pharyngeal endoderm (Fig. 6I, J), pointing to an ancestral role for this transcription factor in pharyngeal endodermal patterning, and possible also in thyroid development.

Our analyses highlighted several genes that were differentially expressed between pharyngeal arch territories, but that were not immediately annotated by BLAST against UniProt/Swiss-Prot, and that required further manual annotation by BLASTing against the larger NCBI non-redundant (nr) database. Among these was *scamp5*, which encodes a secretory carrier membrane protein expressed in the synaptic vesicles of neuroendocrine tissues (Fernández-Chacón & Südhof, 2000; Han et al., 2009), and falls within the same topologically associated domain as single nucleotide polymorphisms associated with orofacial clefting in humans (Carlson et al., 2019). In skate, *scamp5* is a novel marker of dorsal mandibular arch mesenchyme (Fig. 6N, O; Fig. S2G, H). Although *scamp5* has never been previously implicated in pharyngeal arch skeletal patterning, the above observations, combined with our novel *in situ* expression in skate, highlight this gene as a promising candidate for further study. Expression analyses and functional characterisation in bony fish model systems will reveal whether the expression patterns we report here are general features of gnathostomes, or derived features of cartilaginous fishes, and possible undiscovered roles for *scamp5* in craniofacial skeletal development.

### Distinct gene expression features within mandibular and gill arch mesodermal muscle progenitors

The mesodermal cores of vertebrate pharyngeal arches derive from both cranial paraxial and lateral splanchnic mesodermal subpopulations, and give rise to the branchiomeric musculature – i.e. the muscles of mastication and facial expression in mammals, and the muscles of the jaw and gill arches in fishes (Tzahor and Evans, 2011; Ziermann and Diogo, 2019; Sleight and Gillis, 2020). While expression of some elements of the pharyngeal myogenic developmental programme, such as *Tbx1* (Kelly et al., 2004), *Islet-1* (Nathan et al., 2008), *Lhx2* (Harel et al., 2012), myosin heavy chain (Ziermann et al., 2017) and *MyoD* (Schilling and Kimmel, 1997; Poopalasundaram et al., 2019) are shared across the mesodermal cores of multiple pharyngeal arches, other gene expression features are differentially required for the specification of distinct arch-derived muscular features. For example, it has been shown in mouse that *Pitx2* expression within the core mesoderm of the mandibular arch is required for specification of jaw musculature – in part through positive regulation of core mesodermal *Six2* expression – but not for specification of hyoid arch musculature (Shih et al., 2007). It therefore appears as though pharyngeal arch myogenesis is regulated by a core transcriptional programme, with additional arch-specific gene expression directing specific branchiomeric muscle identities.

Our differential expression analyses identified *six2* as enriched in the skate mandibular arch, and *in situ* validation confirmed its expression in the mesodermal core of the mandibular arch at S24 (as well as in the dorsal epithelium of each pharyngeal arch – Fig. 7A, B). We have also identified *tbx18* (Fig. 7C, D) and *pknox2* (Fig. 7E, F) as markers of the mesodermal core of the mandibular arch at S24. *Tbx18* expression within the mandibular arch has previously been reported in mouse (Kraus et al. 2001), zebrafish (Begemann et al., 2002) and chick (Haenig and Kispert, 2004), while *Pknox2* expression has previously been reported from microarray analysis of the mouse mandibular arch (Feng et al., 2009). However, neither *Tbx18* nor *Pknox2* has yet been implicated in the development of mandibular arch-derived musculature. Interestingly, our analyses also revealed *lhx9* as a marker of the mesodermal core of the hyoid and gill arches, but not the mandibular arch (Fig. 7G, H) – a feature so far unreported in any other taxon. Taken together, these findings highlight an ancestral role for *six2* in patterning mandibular arch-derive musculature in jawed vertebrates, possibly in conjunction/parallel with *tbx18* and *pknox2*, as well as *lhx9* as a novel marker of hyoid and gill arch muscle progenitors.

**Figure 7:**
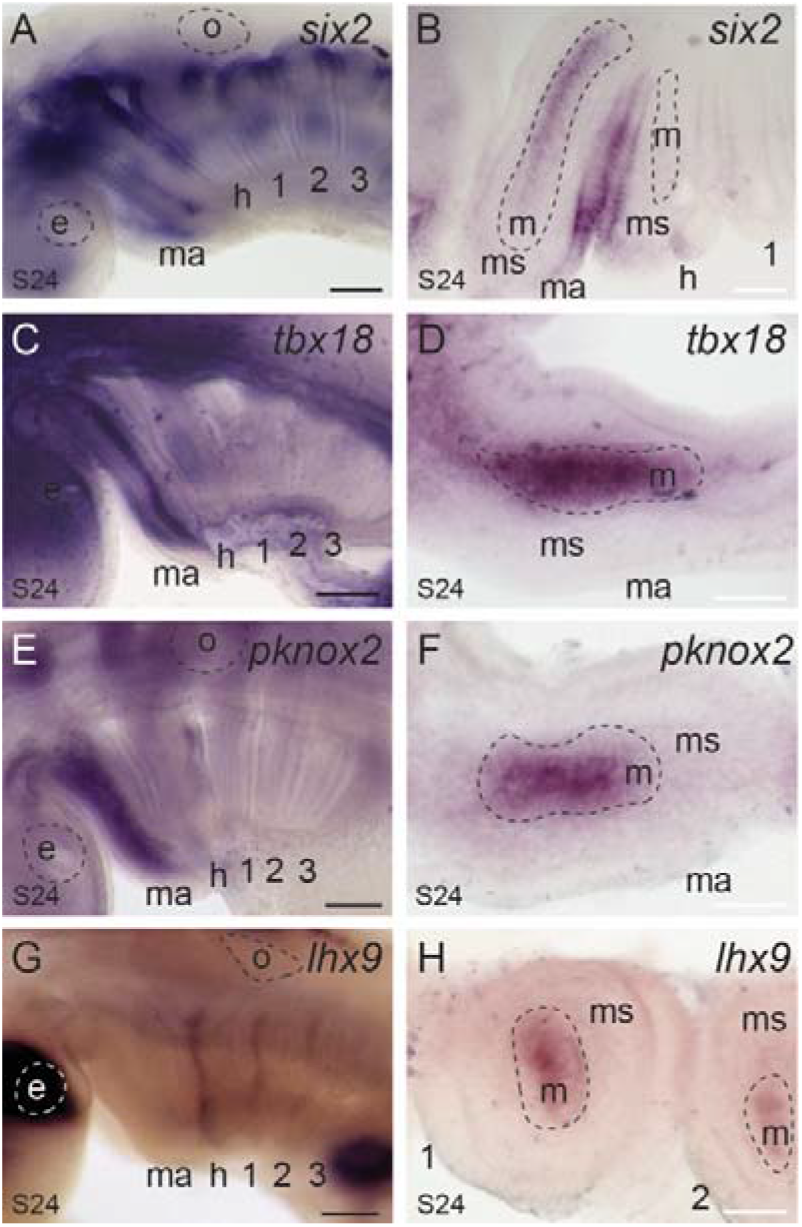
Distinct gene expression features of mandibular and hyoid/gill arch muscle progenitors. (A,B) *six2*, (C,D) *tbx18* and (E,F) *pknox2* are expressed in the core mesoderm of the mandibular arch. Six2 is also expressed in the dorsal epithelium of each pharyngeal arch. (G,H) *lhx9* is expressed in the core mesoderm of the hyoid and gill arches. 1, 2, 3, 4: gill arches 1-4; e, eye; h, hyoid; m, mandibular arch; me, mesoderm; ms, mesenchyme; o, otic vesicle. Scale bars: black 400um, white 25um.

### Gene expression features of presumptive gill epithelium and external gill buds

The gills of fishes derive from the endodermal epithelium of the hyoid and gill arches (Gillis and Tidswell, 2017; Hockman et al., 2017; Warga and Nüsslein-Volhard, 1999). In skate, gills form initially as a series of transient embryonic external gill filaments, which are eventually remodelled and resorbed into internal gill lamellae (Pelster and Bemis, 1992). Our differential expression analysis revealed a number genes to be differentially expressed between the mandibular and first gill arch, some of which proved, through *in situ* validation, to be markers of developing gills. In skate, we observed expression of *foxl2* in the gill-forming endodermal epithelium and developing gill buds of all pharyngeal arches (including the presumptive spiracular pseudobranch primordium – i.e. the precursors of the vestigial gill lamellae of the mandibular arch), as well as in the core mesoderm of each pharyngeal arch (Fig. 8A, B). These expression patterns are consistent with previous reports of *foxL2* expression from mouse (Jeong et al., 2008; Marongiu et al., 2015) and the shark, *Scyliorhinus canicula* (Wotton et al., 2007). We additionally observe expression of *gcm2* throughout the developing gill buds of the hyoid and gill arches (Fig. 8C, D), as well as expression of *wnt2b* (Fig. 8E, F) and *foxQ1* (Fig. 8G, H) in the tips of the developing gill buds. *gcm2* is expressed in the developing gills of shark and zebrafish (Hogan et al., 2004; Okabe and Graham, 2004), and was therefore likely ancestrally required for gill development in gnathostomes. However, there are no previous reports of *wnt2b* or *foxq1* expression during gill development in other taxa, pointing to a possible novel role for these factors in driving outgrowth of external gill filaments.

**Figure 8:**
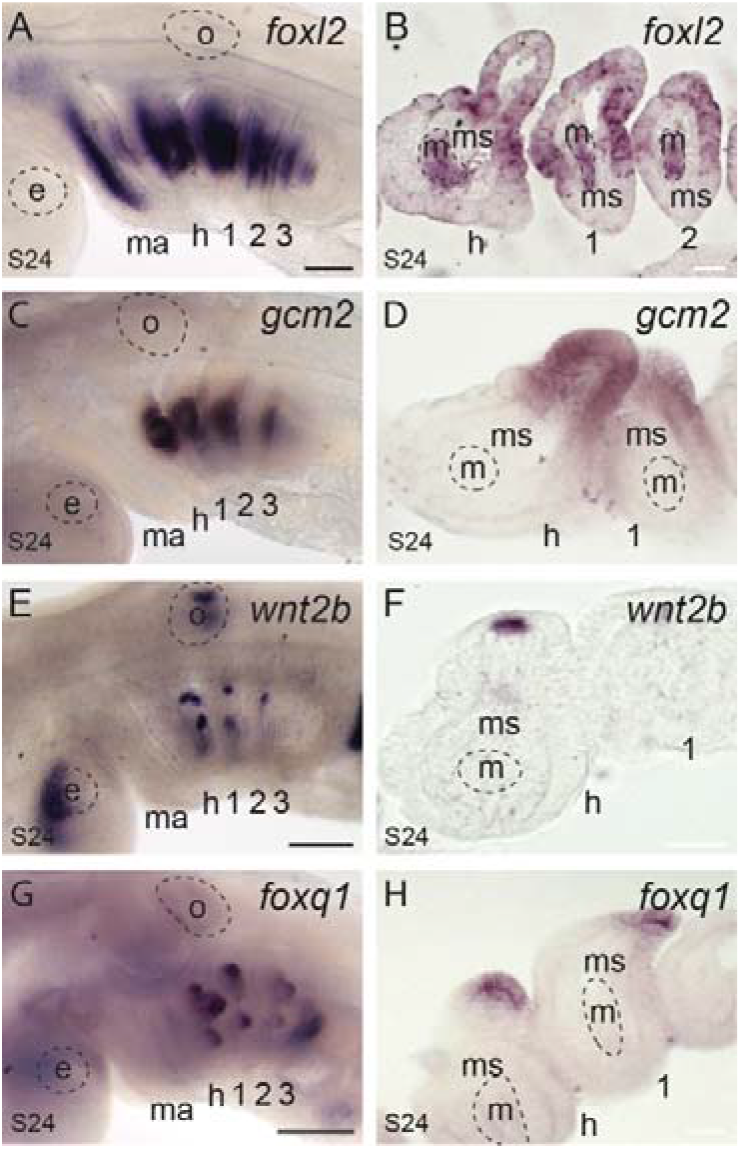
Conserved and novel molecular markers of gill development. (A,B) *foxl2* is expressed in the gill-forming epithelium and core mesoderm of all pharyngeal arches in skate at S24. (C,D) *gcm2* is expressed throughout the developing gill buds of the hyoid and gill arches, while (E,F) *wnt2b* and (G,H) *foxq1* are expressed in the tips of developing gill buds.

### Mandibular and gill arch serial homology and evolution of the jaw

Our combination of candidate and differential gene expression analysis has revealed a suite of transcription and signalling factors that display polarised expression along the DV axis of the pharyngeal arches in skate. The overwhelming majority of genes discussed above share patterns of expression in the mandibular, hyoid and gill arches (Fig. 9A). Together with previous reports of shared expression of core components of the pharyngeal arch DV patterning network in cartilaginous and bony fishes (Gillis et al., 2013; Compagnucci et al., 2013), and the fact that many genes involved in DV patterning of the jaw skeleton in zebrafish have comparable hyoid arch skeletal patterning functions, our findings point to a conserved transcriptional network patterning the DV axis of the mandibular, hyoid and gill arches in the gnathostome crown group, and serial homology of the gnathostome jaw, hyoid and gill arch skeleton. We additionally report distinct transcriptional features of the mandibular and gill arches in skate (Fig. 9B), including dorsal mesenchymal expression of *six1, eya1* and *scamp5*, mandibular arch mesoderm-specific expression of *six2, tbx18* and *pknox2*, hyoid/gill arch mesoderm-specific expression of *lhx9*, and the expression in developing gills of *foxl2, gcm2, wnt2b* and *foxq1*. The aforementioned mesenchymal gene expression features could reflect mandibular arch-specific divergence from the ancestral pharyngeal DV patterning programme, and could function downstream of global anteroposterior patterning mechanisms (e.g. the “Hox code” of the vertebrate head) and in parallel with local signals from oral epithelium to effect anatomical divergence of the mandibular arch skeleton (Hunt et al., 1991; Rijli et al., 1993; Couly et al., 1998, 2002; Hunter and Prince, 2002), while mesodermal and endodermal gene expression features could underlie the evolution of arch-specific muscular and gill fates, respectively.

**Figure 9:**
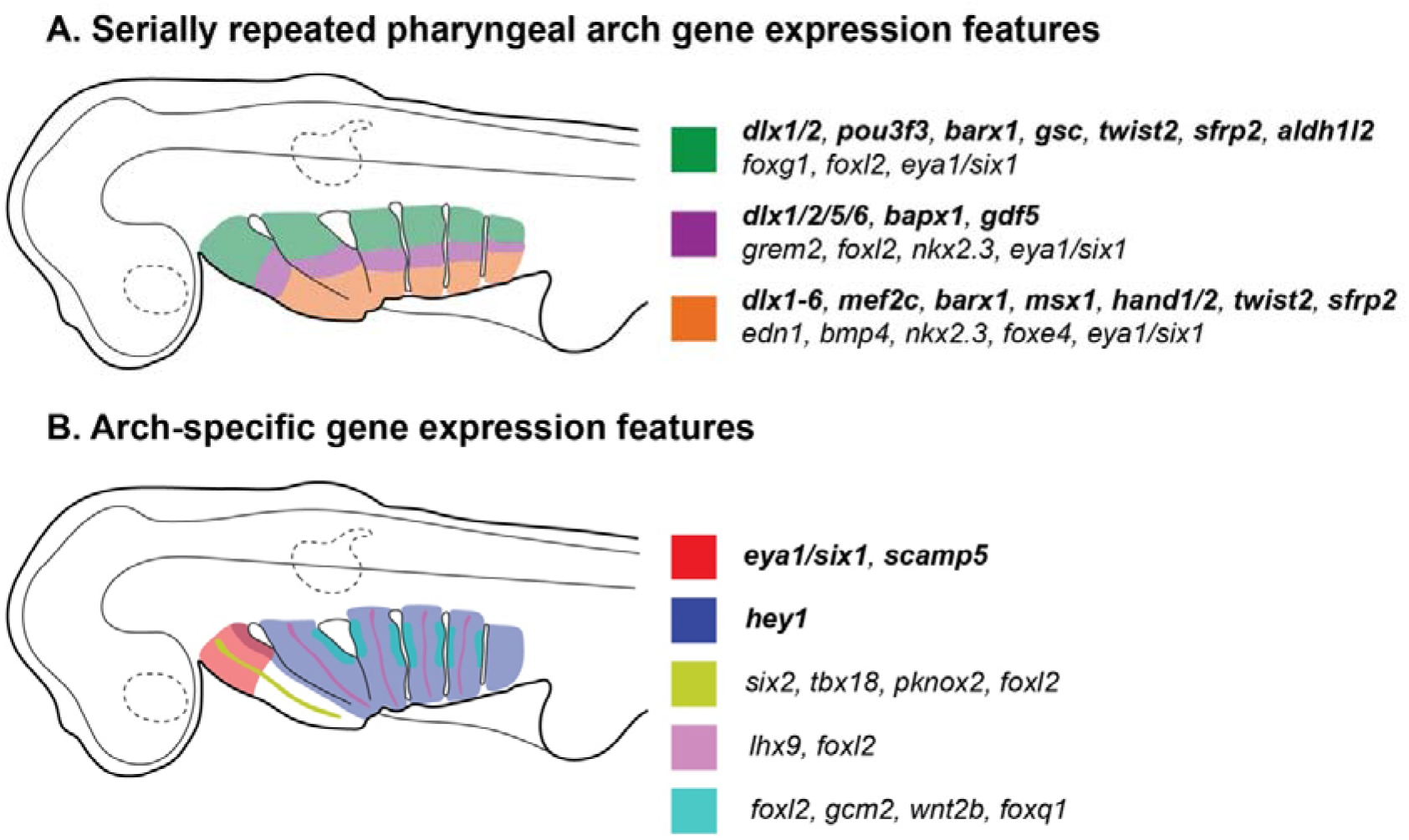
Summary of polarised gene expression patterns within skate pharyngeal arches. (A) Gene expression patterns that are serially repeated across the mandibular, hyoid and gill arches in skate. We propose that these features comprise an ancestral core pharyngeal arch DV patterning programme for gnathostomes, and underlie serial homology of the jaw, hyoid and gill arch skeleton. For schematic purposes, serially repeated gene expression patterns are classified as belonging to one of three broad territories (dorsal, intermediate or ventral). (B) Gene expression features that are unique to one or more pharyngeal arches in skate. Bold italics indicates genes that are expressed in pharyngeal arch mesenchyme, while regular italics indicates genes that are expressed in pharyngeal arch mesoderm and/or epithelium. For details of expression patterns and tissue specificity, please see text.

Cyclostomes (i.e. lampreys and hagfishes) are the most proximate living sister group to the gnathostomes (Heimberg et al., 2010), and the cyclostome pharyngeal endoskeleton and oral apparatus departs considerably from the condition seen in cartilaginous and bony fishes. Lampreys possess a muscular lower lip and lingual and velar cartilages that derive from the first pharyngeal (mandibular) arch, a muscular upper lip that derives largely from the premandibular domain, and a branchial “basket” consisting of a series of unjointed cartilaginous gill, epitrematic and hypotrematic bars, derived from the hyoid and gill arches (Johnels, 1948). This lamprey pharyngeal skeleton arises from embryonic tissue interactions and gene expression patterns that differ from those seen in gnathostomes. For example, lampreys possess six *Dlx* genes of unclear orthology with those of gnathostomes (Neidert et al., 2001; Myojin et al., 2001; Kuraku et al., 2010) – and while these genes are expressed in a nested pattern in the mesenchyme of all pharyngeal arches, this pattern differs from the broadly conserved “Dlx code” that has been described in various gnathostome taxa (Cerny et al., 2010). Additionally, in the rostral pharynx of the lamprey, Dlx-expressing neural crest-derived mesenchyme is not confined to the mandibular arch, but rather extends into the premandibular domain, and patterns of *Dlx* gene expression in this oral region differ from those seen in the posterior pharyngeal arches (Cerny et al., 2010; reviewed by Miyashita and Diogo, 2016).

Most developmental hypotheses of jaw evolution aim to explain the origin of the gnathostome jaw by modification of a cyclostome-like condition – e.g. via a heterotopic shift in epithelial-mesenchymal interactions restricting skeletogenic transcription factor expression to the mandibular arch (Shigetani et al., 2002), by confinement of the embryonic progenitors of ancestrally distinct rostral pharyngeal skeletal elements to the mandibular arch, and subsequent assimilation of mandibular arch derivatives to segmented skeletal arrangement found in more caudal arches (Miyashita, 2015), or by co-option of a developmental mechanism promoting joint fate into a mandibular arch that is otherwise largely gnathostome-like in its DV patterning (Cerny et al., 2010). These hypotheses, in turn, are predicated on the assumption that the cyclostome-like pharyngeal skeleton reflects an ancestral vertebrate condition. There is some palaeontological support for this view, though this comes in the form of inferred cyclostome-like skeletal conditions from casts of cranial nerve paths and muscle scars inside the dermal head shield of stem-gnathostomes, and not from direct observation of endoskeletal preservation (Janvier, 1996).

Preservation of the cartilaginous skeletal elements of early vertebrates is rare, but has been reported for the Cambrian stem-vertebrate *Metaspriggina walcotti* (Conway Morris, 2008), recently reconstructed as possessing seven paired gill bars, each segmented into bipartite dorsal and ventral elements (reminiscent of the epi- and ceratobranchials of crown gnathostomes) (Conway Morris and Caron, 2014). If this reconstruction reflects faithful preservation of the pharyngeal endoskeleton – and if the most rostral of these segmented bars is derived from the first pharyngeal arch – this would imply that a pharyngeal skeletal organization more closely resembling that of crown gnathostomes (i.e. with a serially repeated set of segmented skeletal derivatives arising from each pharyngeal arch) could, in fact, be plesiomorphic for vertebrates. It would follow that differences between cyclostome and gnathostome pharyngeal skeletons reflect cyclostome divergence from the plesiomorphc condition retained in gnathostomes (rather than vice versa), and that the pan-pharyngeal transcriptional programme discussed above could have functioned to pattern the DV axis and to serially delineate pharyngeal skeletal segments not just in the last common ancestor of the gnathostome crown group, but more generally, in the last common ancestor of vertebrates.

## Materials and Methods

### Embryo collection

*L. erinacea* embryos for mRNA *in situ* hybridisation were collected at the Marine Biological Laboratory (Woods Hole, MA, USA). Embryos were fixed in 4% paraformaldehyde overnight at 4°C, rinsed in phosphate-buffered saline (PBS), dehydrated stepwise into 100% methanol and stored in methanol at −20°C. Skate embryos were staged according to Ballard et al. (1993) and Maxwell et al. (2008).

### Gene cloning and mRNA *in situ* hybridisation probe synthesis

Cloned fragments of skate cDNAs were PCR amplified from total embryonic cDNA template using standard protocols. PCR products were isolated and purified using the MinElute Gel Extraction Kit (Qiagen) and ligated into the pGemT-easy Vector System (Promega). Resulting plasmids were transformed into JM109 *E. coli* (Promega), and plasmid minipreps were prepared using the alkaline miniprep protocol by Wang *et al*. (1982). Insert sequences were verified by Sanger Sequencing (University of Cambridge, Dept. of Biochemistry). Linearized plasmid was used as a template for *in vitro* transcription of DIG-labeled riboprobes for mRNA *in situ* hybridization, using 10X DIG-labelled rNTP mix (Roche) and T7 RNA polymerase (Promega), according to manufacturers’ directions. Probe reactions were purified using the RNA Clean & Concentrator kit (Zymo Research).

### Histology and *in situ* hybridization

Paraffin embedding, sectioning and *in situ* hybridizations on sections were performed as described previously (O’Neill et al., 2007; with modifications according to Gillis et al., 2012).

For wholemount *in situ* hybridizations (WMISH), embryos were rehydrated through a methanol gradient into diethylpyrocarbonate (DEPC)-treated phosphate-buffered saline (PBS) with 0.1% Tween-20 (100%, 75%, 50%, 25% methanol in DEPC-PBT), then treated with a 1:2000 dilution of 10mg/mL proteinase K in DEPC PBT for 15 minutes at room temperature. Following a rinse in DEPC-PBT, embryos were re-fixed in 4% PFA/DEPC-PBS for 15 minutes at room temperature and washed in DEPC-PBT again. Specimens were prehybridised in hybridisation solution (5x SSC, 50% formamide, 1% SDS, 50ug/ml yeast tRNA, 25ug/ml heparin) for 1 hour at room temperature. Hybridisation was performed overnight at 70°C with dig-labelled riboprobe diluted to 1ng/uL in hybridisation solution. Embryos were washed twice for 1 hour each at 70°C in wash solution 1 (50% formamide, 2xSSC, 1% SDS), twice for 30 minutes each at 70°C in wash solution 3 (50% formamide, 1xSSC), then three times for 10 minutes at room temperature in MABT (0.1M maleic acid, 150mM NaCl, 0.1% Tween-20, pH 7.5). After blocking for 2 hours at room temperature in 20% sheep serum + 1% Boehringer blocking reagent in MABT, embryos were incubated overnight at 4°C with a 1:2000 dilution of anti-digoxigenin antibody (Roche) in blocking buffer. Embryos were then washed in MABT (two quick rinses then five 30-minute washes), stored overnight in MABT at 4°C and equilibrated in NTMT (100mM NaCl, 100mM Tris pH 9.5, 50mM MgCl2, 0.1% Tween-20). The colour reaction was initiated by adding BM Purple (Merck) to the embryos, and stopped by transferring to PBS. Embryos were rinsed once in PBS, post-fixed in 4% PFA for 30 minutes, and graded into 75% glycerol in PBS for imaging.

For gelatin embedding, WMISH embryos were equilibrated in a 15% w/v gelatin solution in PBS at 50°C for 1 hour, before being poured into plastic moulds, positioned for sectioning and left to cool. Gelatin block were then post-fixed in 4% PFA at 4°C for 4 days and rinsed in PBS. 50μm sections were cut using a Leica VTS1000 vibratome and mounted on Superfrost slides (VWR) using Fluoromount G (SouthernBiotech).

### RNAseq, *de novo* transcriptome assembly, and differential gene expression analysis

Total RNA was extracted from upper mandibular arch (n=10), lower mandibular arch (n=6), upper gill arch (n=5) and lower gill arch (n=3) domains at stage (S)23/24 and from upper mandibular arch (n=6), lower mandibular arch (n=6), upper gill arch (n=4) and lower gill arch (n=4) domains at S25/26 (Fig. 5A). S23-24 and S25-26 span the expression of the *dlx* code, a key regulator of axial identity in the pharyngeal arches. Unlike in mouse and zebrafish, where combinatorial *dlx* expression has only been observed in the mandibular and hyoid (Depew et al., 2002; Talbot et al., 2010), the typically nested expression of the *dlx* code is shown by the developing mandibular, hyoid and gill arches in skate (Gillis et al., 2013). Manual dissections of upper and lower arch primordia were guided by morphological landmarks correlating with dorsal (*dlx1/2+*) and ventral (*dlx1-6*+) expression (after Gillis et al., 2013).

Samples were preserved in RNAlater, total RNA was extracted using the RNAqueous-Micro Total RNA Isolation Kit (ThermoFisher), and library prep was performed using the Smart-seq2 (Picelli et al., 2014) with 10 cycles of cDNA amplification. S23/24 and S25/26 libraries were pooled and sequenced using the HiSeq4000 platform (paired end sequencing, 150bp read length) at the CRUK genomics core facility (University of Cambridge, Cancer Research UK Cambridge Institute). In addition to the above, libraries from the dorsal mandibular arch (n=5), ventral mandibular arch (n=5), dorsal gill arch (n=5) and ventral gill arch (n=5) domains of S29 skate embryos were prepared as described above, and sequenced using the NovaSeq 6000 (paired end sequencing, 150bp read length) at Novogene Co., Ltd. Reads from these libraries were included in our *de novo* transcriptome assembly, but are not analysed further in the current work.

A total of 2,058,512,932 paired raw reads were used. Low quality read and adapter trimming was conducted with Trim Galore! (<u>0.4.4</u>) with the quality parameter set to 30 and phred cut-off set to 33. Reads shorter than 65 bp were discarded. After trimming adapters and removing low quality reads a total of 1,348,098,076 reads were retained. Normalisation (max coverage 30) reduced this to a further 54,346,196 reads. The *de novo* strand-specific assembly based on these reads was generated using Trinity 2.6.6 with default parameters (Grabherr et al., 2011; Haas et al., 2013). The N50 is 1009bp, and the Ex90N50 (the N50 statistic computed as usual but considering only the topmost highly expressed transcripts that represent 90% of the total normalized expression data, meaning the most lowly-expressed transcripts are excluded) is 1906bp (Table S1). Post-assembly quality control was carried out using Trinity’s toolkit or gVolante (Fig. S2., Table S1).

Trinity transcript quantification was performed alignment-free using salmon (Patro et al., 2017) to estimate transcript abundance in TPM (Transcripts Per Kilobase Million). The genes differentially expressed along the DV axis within each arch, or across the anterior-posterior axis between dorsal and ventral elements of each arch, were screened for using edgeR with a cutoff of FDR (false discovery rate) ≤0.05 (Table S2 for gene numbers, Table S3 for candidates for validation, Tables S4-7 for stages 24/25 and Tables S8-11 for stages 25/26). Within EdgeR, a negative binomial additive general linear model with a quasi-likelihood F-test was performed. Model design accounted for repeated sampling of tissues from the same individual and p-values were adjusted for multiple testing using the Benjamin-Hochberd method to control the false discovery rate (FDR ≤0.05) (Fig. S1 A-D). The transcripts were putatively annotated based on sequence similarity searched with blastx against Uniprot (http://www.uniprot.org/).

## Acknowledgements and funding information

We thank Dr. Richard Schneider, Prof. David Sherwood and the MBL Embryology course for provision of lab space, Louise Bertrand and Leica Microsystems for microscopy support and David Remsen, Scott Bennett, Dan Calzarette and the staff of the Marine Biological Laboratory and MBL Marine Resources Center for technical and animal husbandry assistance. We also thank Jenaid Rees, Dr. Kate Rawlinson and the University of Cambridge chordate evo-devo community for helpful discussion.

*Leucoraja erinacea* sequence data from this work are available under the following NCBI accession numbers: SRA data (PRJNA686126) and accession numbers MN478367, MW389327-MW389328 and MW457797-MW457625.

This work was supported by a Biotechnology and Biological Sciences Research Council Doctoral Training Partnership studentship to CH, by a Wolfson College Junior Research Fellowship and MBL Whitman Early Career Fellowship to VAS, and by a Royal Society University Research Fellowship (UF130182 and URF\R\191007), Royal Society Research Grant (RG140377) and University of Cambridge Sir Isaac Newton Trust Grant (14.23z) to JAG.

## Supplemental Material

**Supplemental Table 1:**
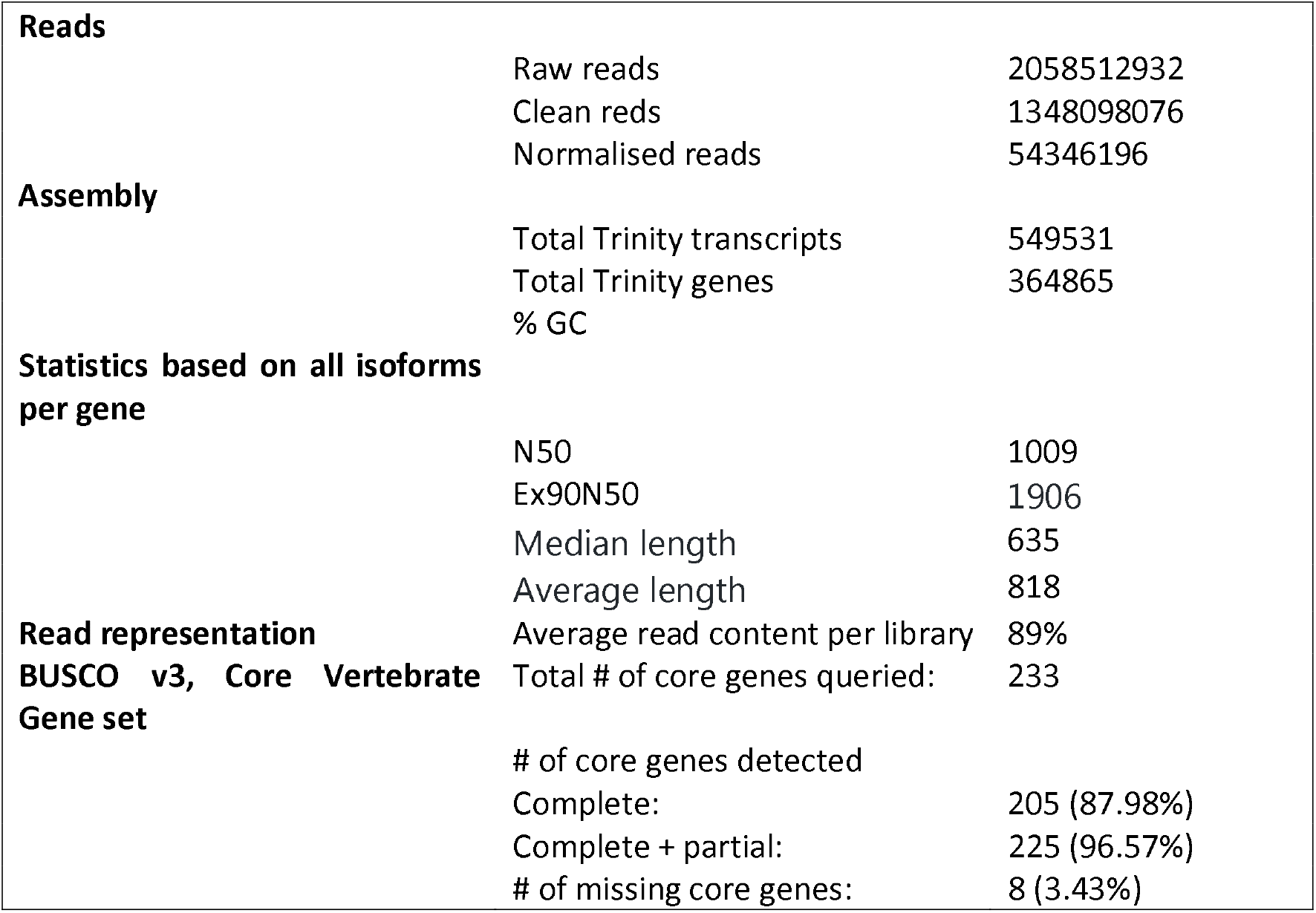
Skate pharyngeal arch transcriptome assembly statistics.

**Supplemental Table 2:** Number of significantly expressed transcripts per differential gene expression analysis between and within arches across the DV axis.

|  | Gill arch<br>S23_24 | Mandibular Arch<br>S23_24 | Gill Arch<br>S25_26 | Mandibular Arch<br>S25_26 |
| --- | --- | --- | --- | --- |
| Dorsal | 10 | 57 | 11 | 46 |
| Ventral | 9 | 54 | 21 | 44 |
|  | Dorsal S23_24 | Ventral S23_24 | Dorsal S25_26 | Ventral S25_26 |
| Gill arch | 238 | 83 | 45 | 919 |
| Mandibular arch | 200 | 105 | 71 | 691 |

**[Supplemental Tables 3-11 provided as Excel spreadsheets]**

**Supplemental Table 3:** Candidate genes differentially expressed in skate pharyngeal arches considered for further validation by mRNA *in situ* hybridisation.

**Supplemental Figure 1:**
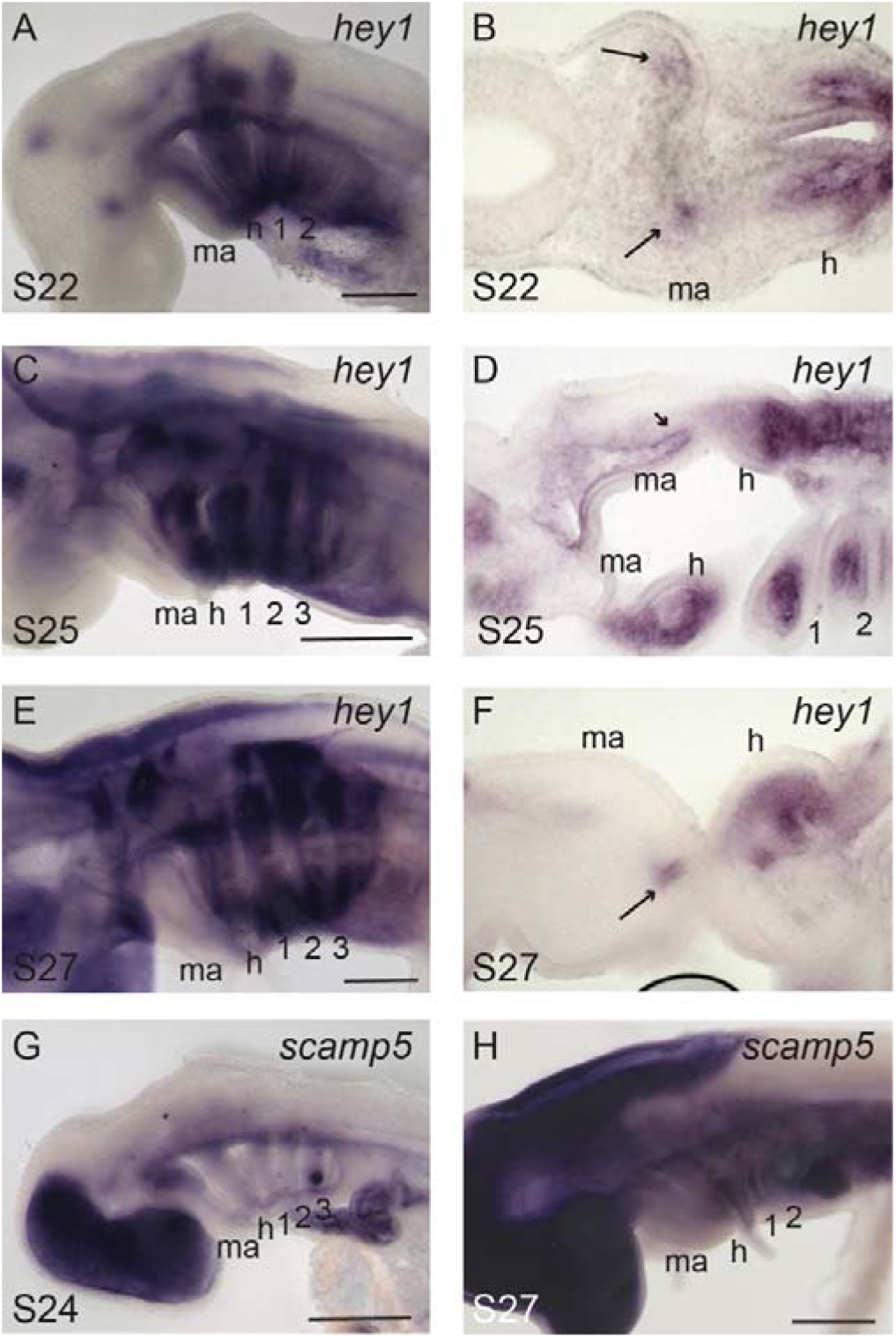
Additional expression of *hey1* and *scamp5* in skate pharyngeal arches. At (A,B) S22, (C,D) S25 and (E,F) S27, *hey1* is highly expressed in the mesenchyme of the hyoid and gill arches, but expressed only in a very restricted domain in posterior mandibular arch mesenchyme (arrows in B, D and F). (G) *scamp5* expression in dorsal mandibular arch mesenchyme persists in S24, but (H) is no longer detectable at S27. 1, 2, 3, 4: gill arches 1-4; h, hyoid; m, mandibular arch; Scale bars: black 1mm, white 25um.

**Supplemental Figure 2:**
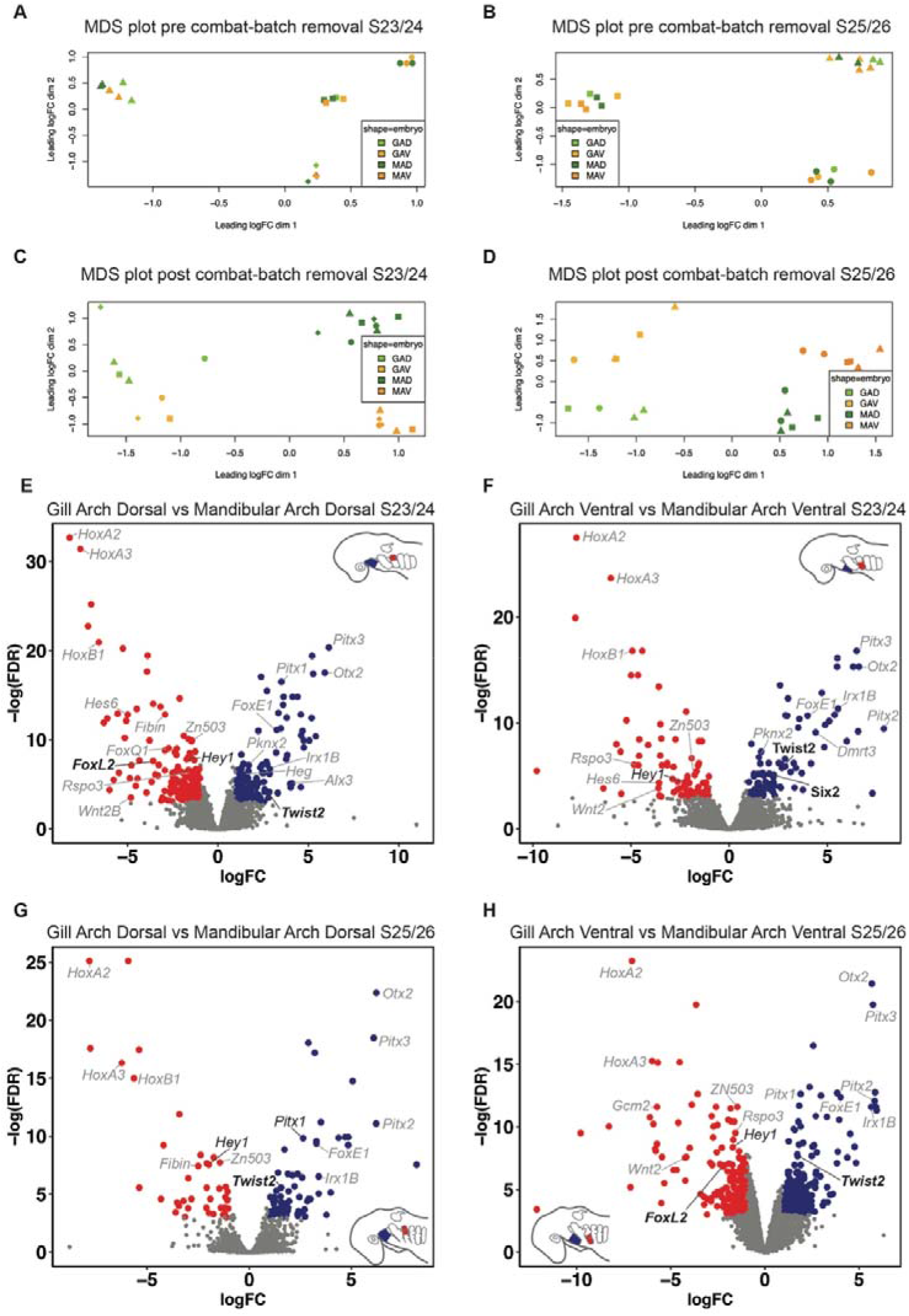
MDS plots revealing sample heterogeneity and Volcano plots showing genes differentially expressed in skate pharyngeal arches. (A) Multi-dimensional scaling (MDS) plots show samples both in S23/24 and (B) S25/26 cluster more according to embryo of origin than tissue type (GAD = gill arch dorsal, GAV= gill arch ventral, MAD = mandibular arch dorsal, MAV = mandibular arch ventral). (C-D) This batch effect was accounted for during statistical testing. After combat adjustment using empirical Bayes methodology (Johnson et al., 2007), MDS plots show samples clustering according to tissue type. Volcano plots illustrate genes that are significantly differentially expressed (E) between the dorsal domains of the mandibular and the gill arch at S23/24, (F) between the ventral domains of the mandibular and the gill arch at S23/24, (G) between the dorsal domains of the mandibular and the gill arch at S25/26 and (H) between the ventral domains of the mandibular and the gill arch at S25/6. Genes with established roles in pharyngeal arch axial patterning are in simple italics, additional genes for which we provide *in situ* validation are in bold italics, and additional factors highlighted by our analysis but not validated by mRNA *in situ* hybridisation are in grey italics.

## Notes

### Competing Interest Statement

The authors have declared no competing interest.

